# Retinal thermometry in-vivo using phase-sensitive optical coherence tomography

**DOI:** 10.1101/2024.08.07.607046

**Authors:** Yueming Zhuo, Mohajeet Bhuckory, Huakun Li, Junya Hattori, Davis Pham-Howard, David Veysset, Tong Ling, Daniel Palanker

## Abstract

Controlling the tissue temperature rise during retinal laser therapy is essential for predictable outcomes, especially at non-damaging settings. We demonstrate a method for determining the temperature rise in the retina using phase-resolved optical coherence tomography (pOCT) in vivo. Measurements based on the thermally induced optical path length changes (ΔOPL) in the retina during a 10-ms laser pulse allow detection of the temperature rise with a precision less than 1 °C, which is sufficient for calibration of the laser power for patient-specific non-damaging therapy. We observed a significant difference in confinement of the retinal deformations between the normal and the degenerate retina: in wild-type rats, thermal deformations are localized between the retinal pigment epithelium (RPE) and the photoreceptors’ inner segments (IS), as opposed to a deep penetration of the deformations into the inner retinal layers in the degenerate retina. This implies the presence of a structural component within healthy photoreceptors that dampens the tissue expansion induced by the laser heating of the RPE and pigmented choroid. We hypothesize that the thin and soft cilium connecting the inner and outer segments (IS, OS) of photoreceptors may absorb the deformations of the OS and thereby preclude the tissue expansion further inward. Striking difference in the confinement of the retinal deformations induced by a laser pulse in healthy and degenerate retina may be used as a biomechanical diagnostic tool for the characterization of photoreceptor degeneration.

## Introduction

Precise control of the temperature rise in tissue during laser therapy is essential for predictable and reproducible outcomes. The energy absorption and associated temperature rise depend on the irradiation parameters (wavelength, power, duration, spot size, etc.) and on tissue properties (light scattering and absorption, thermal conductivity, heat capacity, etc.). In retinal laser treatments (1), light delivered to the retina is absorbed primarily by melanin, which is highly concentrated in the retinal pigmented epithelium (RPE) and pigmented choroid (2, 3). Melanin concentration varies significantly between individuals, as well as across the retina within the same subject, up to a factor of four (4). Moreover, the transmittance of the transparent ocular tissues (cornea, lens, vitreous and inner retina) decreases with age and might be affected by diseases, hence reducing the temperature rise in the retina (5, 6). If such variations are not accounted for, they can result in over- or under-treatment of the retina.

The dosimetry of hyperthermia can be quantified assuming that cells respond to a decrease in concentration of biomolecules due to thermal denaturation. Based on the Arrhenius equation and assuming first-order reaction kinetics (7), the integration of the reaction rate over time yields the total fraction of the denatured biomolecules during hyperthermia. This value is further normalized to Ω = 1 to define the threshold of tissue damage. It has been established that retinal response to hyperthermia becomes noticeable when Ω *>* 0.1, thereby defining the therapeutic window of nondamaging thermal therapy to 0.1 *<* Ω*<* 1 (8–10).

In current clinical practice, retinal laser damage threshold is often determined by subjective evaluation of the test lesions in the periphery of the retina by a physician (11). Recently, a photoacoustic system was introduced, which uses a pulsed laser to induce thermoelastic expansion of melanosomes, and measures the pressure transients to determine the temperature and optimize the laser power or exposure duration for a desired treatment outcome (12). This technique was not yet accepted in clinical practice due to the need for a more expensive and complex laser system and photoacoustic detection. Mapping of the retinal temperature rise in response to low-power laser pulses by phase-sensitive optical coherence tomography (pOCT) prior to laser treatment has been proposed as a non-invasive alternative (13, 14). In the future, when OCT machines will be upgraded to phase-sensitive detection, such mapping could be performed by the same tool as the structural OCT, and laser settings could be adjusted automatically during treatment according to these maps using systems such as PASCAL or Navilas (15, 16). Optical path length changes (ΔOPL) in OCT provide high sensitivity in detecting minute thermal responses in tissue, and have been shown to correlate with temperature measurements (17, 18). In our previous work, we have demonstrated how pOCT allows for precise detection of laser-induced thermal transients exvivo (19). Tissue parameters and temperature profiles were extracted using a thermo-mechanical modeling (20), which was subsequently validated by a temperature-sensitive fluorescence imaging (19).

In this study, we demonstrate that a similar approach can be reproduced in vivo, utilizing a subpixel registration algorithm for compensating bulk tissue motion due to heartbeat, breathing cycle, and eye movements. Our results reveal an unexpected difference in the thermal deformation patterns between normal and degenerate retina: the OPL changes in healthy retina are confined within the photoreceptor outer segments, as opposed to a much deeper penetration of the tissue expansion in the degenerate retina. These findings indicate the presence of a soft structural element between the photoreceptors’ outer segments (OS) and inner segments (IS), which we hypothesize to be the interconnecting cilium, that dampens the thermal expansion of the OS driven by laser heating of underlying pigmented layers: RPE and choroid. We tested the validity of this hypothesis by thermomechanical modeling. This phenomenon may provide diagnostic insight in evaluating the health of photoreceptors.

## Results and Discussion

### Measuring the thermally induced OPL change with pOCT

Animal preparations and the retinal heating experiments were conducted under room light conditions. Two strands of pigmented rats have been used: wild-type Long Evans (LE) rats and Royal College of Surgeons (RCS) rats having retinal degeneration. To prevent optoretinographic (ORG) responses (21), the wild-type (WT) rat retinas were fully bleached using a full-field illumination from a green LED (CW, 50 µW, centered at 565 nm, illumination area about 6 mm^2^, bleached for 3 minutes). RCS rats exhibit photoreceptor degeneration due to RPE’s inability to phagocytose rod outer segments (22), and their light-insensitive retinas were used to validate that thermal deformations are not induced by ORG responses. Animals were imaged by a custom-built high-speed line-scan spectral-domain optical coherence tomography (LS-SD-OCT) system (see *Materials and Methods*). Retinal cross-sectional scans were captured at 10 kHz frame rate. Heating laser beam at 532 nm wavelength was applied for 10 ms at 10.8 mW of power measured in front of the eye. To minimize motion caused by heartbeat and breathing, rat’s head was stabilized using a custom-built stage equipped with a bite bar and ear bars.

An averaged B-scan of a 6-month-old WT rat is shown in Fig. 1A. The labeled IS/OS line is commonly attributed to the ellipsoid zone (EZ) of the inner segments, which is rich in mitochondria (23, 24). The connecting cilium, a narrow bridge linking the inner and outer segments of photoreceptors, is also suggested to be part of the hyperreflective IS/OS band (25). This structure is crucial for the transport of proteins and other molecules necessary for the function and maintenance of the photoreceptor outer segments (26). The thick hyperreflective band between the IS/OS junction and the Bruch’s membrane (BrM) is composed of the rod outer segments and RPE, including the RPE microvilli interdigitating with the rod outer segment tips (ROST) (27). As shown in the averaged B-scan (Fig. 1B) of the same age RCS rat retina, the degenerated rod outer segments have formed a hyperreflective layer (HRL) above the RPE. The imaging quality of the WT retina is generally better than RCS retina due to healthier eye optics.

**Fig. 1.**
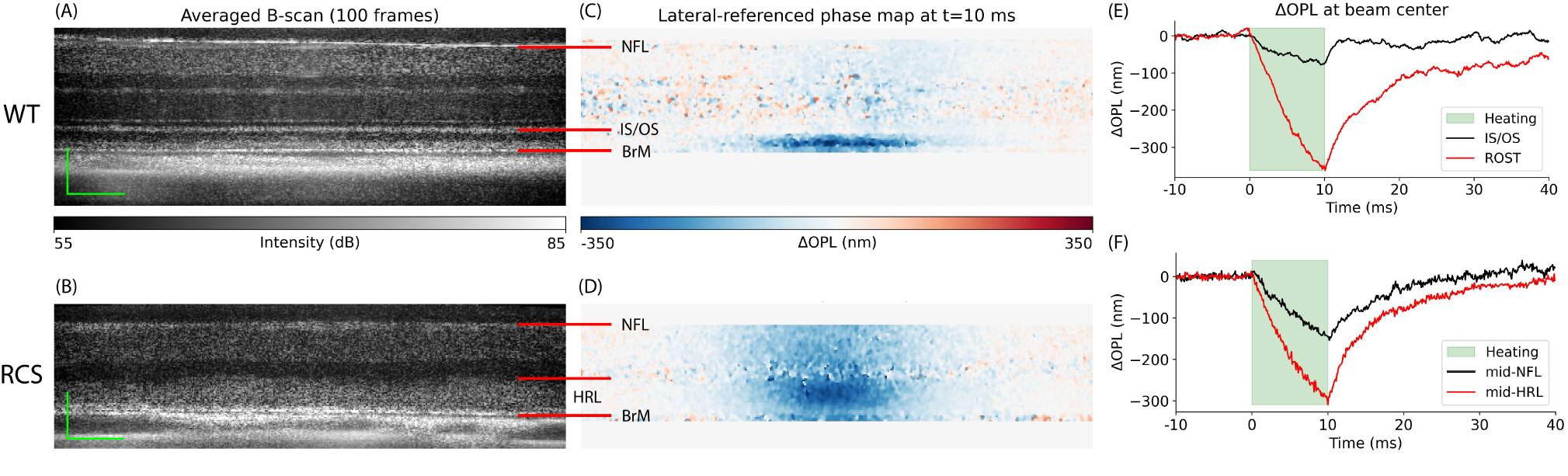
Phase-sensitive OCT imaging of the retinal deformations. (A) Averaged B-scan of a 6-month old wild-type (WT) rat retina. SNR at BrM: 30 dB. Scale bar: axial - 90 μm; lateral - 50 μm. (B) Averaged B-scan of a 6-month old RCS rat retina. SNR at RPE: 30 dB. Scale bar: axial - 70 μm; lateral - 50 μm. (C-D) Laterally referenced phase maps for WT and RCS rats at t = 10 ms. (E-F) ΔOPL change over time at beam center for WT and RCS rats. NFL: nerve fiber layer. IS/OS: inner segment and outer segment. BrM: Bruch’s membrane.

To correct for bulk tissue motion between repeated B-scans, a custom image registration algorithm, called phase-restoring subpixel motion correction (PRSMC) was employed (28). To map the tissue deformations induced by heating, we calculated pixelwise phase changes between subsequent frames and the first frame. To eliminate the residual phase changes induced by the rigid motion of the retina, such as eye rotation, edges of the OCT B-scan were used as the lateral reference region (see *Materials and Methods* and *Supplementary Information*, Fig. S3). Doing so, we obtained the tissue dynamics with respect to the edges of the OCT B-scan (Figs. 1C-D). As expected, during the laser heating (Figs. 1E-F), thermal expansion of tissue leads to a negative change in OPL from the combined effects of tissue moving anteriorly and decrease in refractive index as the temperature rises. After the laser pulse, heat diffuses away and tissue cools down, gradually returning to its initial state.

A significant difference was observed between the ΔOPL maps of RCS and WT retinas. In RCS retina, the OPL change decreases from its maximal value in the HRL to about half of it in the NFL (Figs. 1D, F). However, in WT retina, the OPL changes of a similar magnitude are consistently confined between BrM and IS/OS junction, with negligible changes above it (see Fig. S4 in *Supplementary Information* for repeatable signal patterns from different WT rats). This pronounced localization implies that the WT retina has a particularly soft structural element within the photoreceptor layer that effectively absorbs the thermal strain originating from the heating of RPE and pigmented choroid layers below.

### Thermo-mechanical modeling of the retina

Thermally induced OPL changes can be attributed to tissue expansion and to changes in refractive index. To extract the tissue temperature, a model of ΔOPL should be computed, based on the temperature fields and the vertical displacements of retinal layers, which are both solutions to a thermo-mechanical problem with coupled equations of elasticity and heat diffusion. Here, we briefly describe the model setup, following (19, 20).

The retina is modeled as an axisymmetric multi-layered medium composed of isotropic elastic solid layers. Thermal stress arises from the heat generated by the laser energy absorbed in the RPE and in pigmented choroid. We define the optical transmittance through the eye to the RPE layer as *η*, the spatio-temporal profile of the heat flux at the RPE layer along the depth *z* can be written based on the Beer-Lambert law:

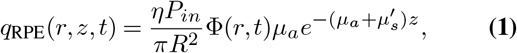

where *P*_*in*_ denotes the input power measured in front of the eye, *R* is the heating spot radius, Φ (*r, t*) is the spatiotemporal profile of the laser pulse with *r* and *t* denoting radial and temporal components respectively, *µ*_*a*_ is the absorption coefficient, and 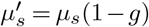 is the reduced scattering coefficient with *μ*_*s*_ being the scattering coefficient and *g* the anisotropy factor. The local heat flux at the pigmented choroid layer can be written in a similar fashion, taking into account that the incident laser power is attenuated by the RPE layer above.

For modeling purposes, all volumetric heat sources were discretized into surface heat sources that are sufficiently close to each other, so the characteristic heat diffusion time across them is much shorter than the 10-ms laser pulse duration (19). For instance, heat diffusion time across 1-µm thick RPE apical layer is about 2.5 µs. Magnitude of the surface heat source at the top of the RPE apical layer was set to be equal to the total heat deposited in it (see *Supplementary Information*). The 20-µm thick pigmented choroid layer was discretized into five 4-µm thick layers, and a surface heat source was applied at the top of each layer.

In cylindrical coordinates, assuming quasi-static conditions, the governing partial differential equations of motion in equilibrium for an elastic layer are:

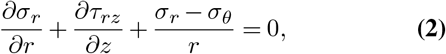

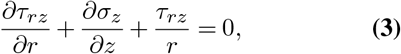

where *σ*_*r*_, *σ* _*θ*_, and *σ*_*z*_ are the normal stress components, *τ*_*rz*_ is the shear stress in the *r − z* plane. The isotropic stress-strain constitutive relations including the thermal stress are:

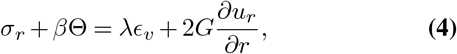

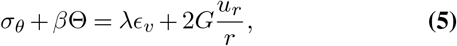

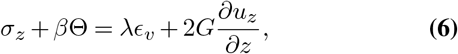

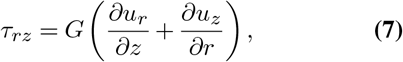

where *u* denotes displacements, *G* is the shear modulus, *λ* is the Lamé’s first parameter, ϵ_*v*_ denote the volumetric strain, and Θ denotes the temperature rise. The thermo-mechanical coupling parameter *β* is linear to the thermal expansion coefficient *α*_TE_, i.e.,

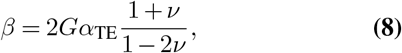

where *v* is the Poisson’s ratio. The heat transfer is described by the Fourier’s heat conduction law:

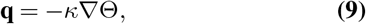

and the heat diffusion equation:

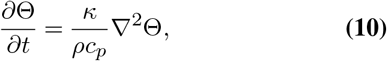

where **q** = [*q*_*r*_, *q*_*z*_]^T^ is the heat flux vector, ∇ and ∇^2^ denote gradient operator and Laplacian operator, respectively. *κ* is the thermal conductivity, *ρ* is the material density, and *c*_*p*_ is the material specific heat capacity. The heat flow in the *z*direction can be computed by:

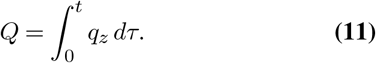

All above equations are coupled to form the complete thermo-mechanical problem of a single layer.

The solution to the thermo-mechanical problem of a multilayered medium with several absorptive layers is fully derived in the *Supplementary Information*. In brief, by transforming the above equations into the Hankel-Laplace (HL) domain, i.e., transforming from the primal domain (*r, z, t*) to HL domain (ξ, *z, s*), a stress vector 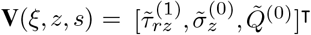 is related to a state vector 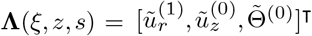 by a transfer matrix **K**(ξ, *z, s*) for a single elastic layer (29–32), i.e.,

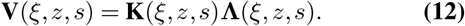

The superscript denotes the order of HL transform. Matrix equations at different depths can be arranged together for the multi-layered retina model. The state vector can be solved by simple matrix multiplication. The numerical inverse HL transform converts the solution back into the primal domain.

Retinal temperature derived from this analytical model depends on the assumed biomechanical properties. Various methods have been employed to measure mechanical properties of ocular tissues, including atomic force microscopy (AFM) (33, 34) and optical coherence elastography (OCE) (35–37). In this study, we use the biomechanical properties listed in Table S1 (see *Supplementary Information*), which are based on values reported in the literature and determined in our previous ex-vivo studies (19).

Temperature rise and displacement fields in thermomechanical model of the WT retina are shown in Fig. 2. ΔOPL can be calculated at any point in space and time based on the temperature and tissue displacements (Fig. 2D, *Materials and Methods*). To account for the strongly confined OPL change observed in Fig. 1C, we added a soft elastic layer, corresponding the connecting cilium layer (CCL), between the IS/OS line and the rod outer segments. The connecting cilium (CC) is a thin, 0.3-0.5 µm wide, 1-2 µm long structure that forms the only physical connection between the IS and OS (26, 38). Surrounding the CC areas are relatively devoid of cellular structures, often referred to as the periciliary membrane space (39, 40). Vertical placement of this junction varies between cells, forming a thicker hyperreflective line in the OCT B-scan. Note that addition of this layer does not change the temperature field (Fig. 2A), but the displacement field becomes drastically different (Figs. 2B-C).

**Fig. 2.**
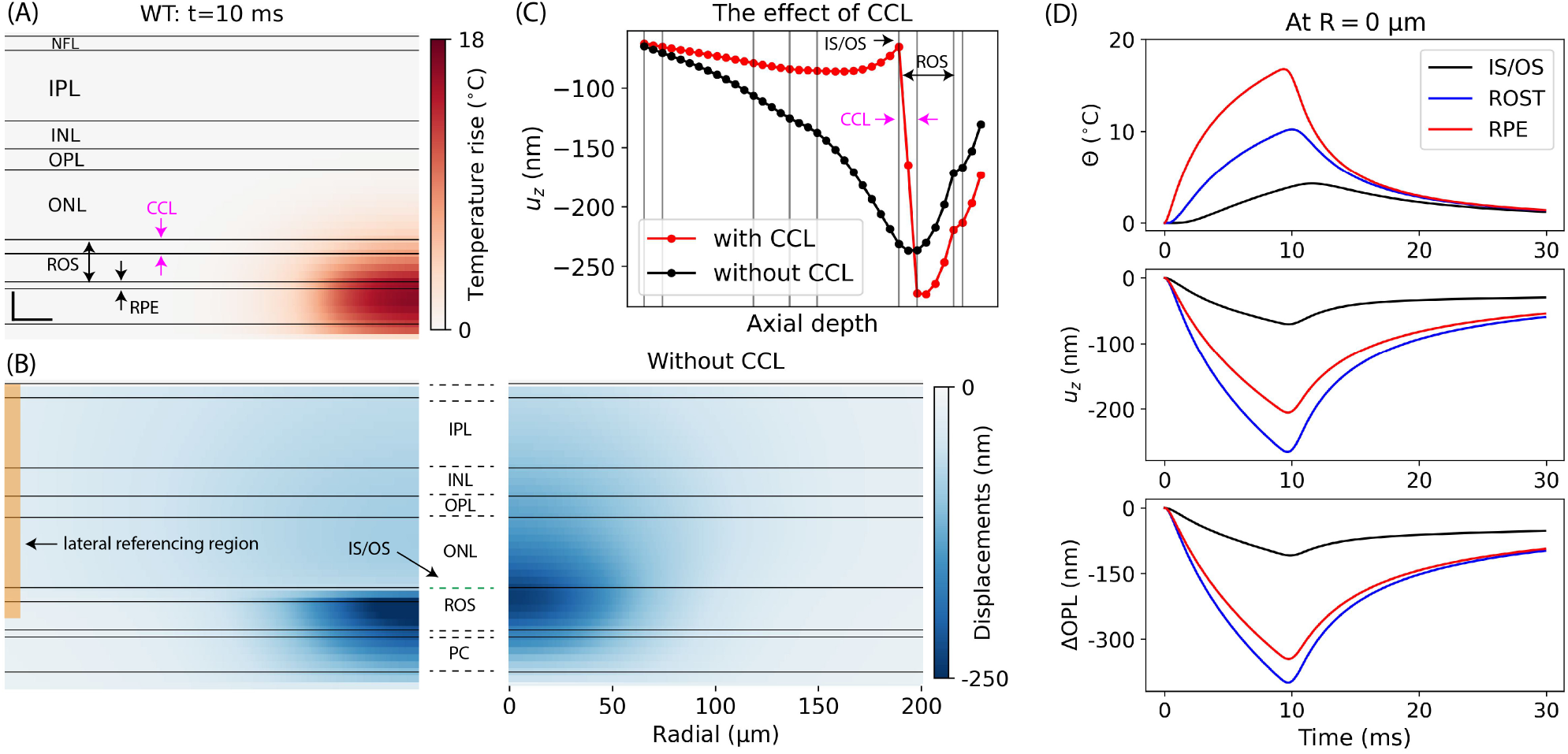
Thermo-mechanical simulation results with parameters: *η* = 50%, *C*_*μ*_ = 0.5, *R* = 45 µm. (A) Simulated temperature map at *t* = 10 ms. Scale bar: axial - 20 μm; lateral - 20 μm. (B) Simulated displacement map at *t* = 10 ms with the CCL layer (left) and without it (right). To determine the reference OPL change similarly to the experimental phase processing (see *Materials and Methods*), the OPL changes within the lateral referencing region were averaged (yellow rectangle). (C) Tissue displacement profile showing the effect of the CCL layer. (D) Time course of displacements, temperature, and OPL change at the beam center for various layers. CCL: connecting cilium layer.

Thickness and stiffness (Young’s modulus) of this soft layer are strongly correlated to maintain the same level of mechanical damping. With its thickness assumed to be equal to that of IS/OS line (about 10 µm), to match the OPL change above the IS/OS line up to NFL to experimental values (Fig. 1C), we adjusted the Young’s modulus to 20 Pa. Such a low Young’s modulus was reported in some neuronal cells (41, 42). Thinner layer would have to be even softer to provide a similar extent of mechanical damping.

### Fitting the model parameters

The model fitting parameters include the ocular optical transmittance, absorption coefficient (proportional to the local melanin concentration) and eye magnification, which affects the heating laser spot size. Denoting a scaling factor *C*_*µ*_ relative to the literature value of the absorption coefficient, i.e., 9.976 *×* 10_3_ cm^*−*1^ for RPE and 2.494 *×* 10_3_ cm^*−*1^ for the pigmented choroid layer (43), the list of the fitting parameters chosen for this study is: *η, C*_*µ*_, and *R*. Note that, *C*_*µ*_ scales the absorption coefficient of both the RPE layer and the pigmented choroid layer. The tissue thermal expansion coefficient was taken from our previous work (19) and all other tissue properties were listed in Table S1. The initial heating laser radius is chosen based on the geometrical optics approximation of the rat eye (44). The optimization objective is to minimize the squared Frobenius norm of the difference matrix between the experimental and model-derived ΔOPL data:

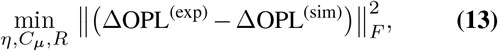

where the subscript *F* denotes the Frobenius norm. The plane near the upper boundary of the ROST-RPE complex was extracted as the ΔOPL^(exp)^ (shown as the blue line in Fig. 3A). The distance between the center of the IS/OS line and the selected plane was measured via the structural OCT image and set in the model such that the ΔOPL^(sim)^ plane is the same distance away from the IS/OS line. Good agreement was obtained between the model and the experimental ΔOPL data both radially and temporally (Figs. 3B-C). The best-fit parameters and the peak temperature predictions are summarized in Table 1.

**Table 1.**
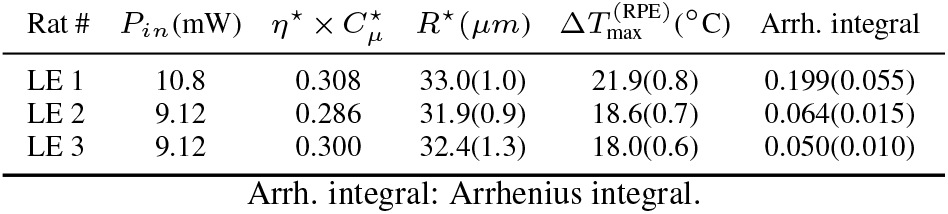
Parameter fitting and temperature estimation results.

| Rat # | $P_{in}$(mW) | $\eta^* \times C_\mu^*$ | $R^*$(μm) | $\Delta T_{max}^{(RPE)}$(°C) | Arrh. integral |
| --- | --- | --- | --- | --- | --- |
| LE 1 | 10.8 | 0.308 | 33.0(1.0) | 21.9(0.8) | 0.199(0.055) |
| LE 2 | 9.12 | 0.286 | 31.9(0.9) | 18.6(0.7) | 0.064(0.015) |
| LE 3 | 9.12 | 0.300 | 32.4(1.3) | 18.0(0.6) | 0.050(0.010) |

Arrh. integral: Arrhenius integral.

**Fig. 3.**
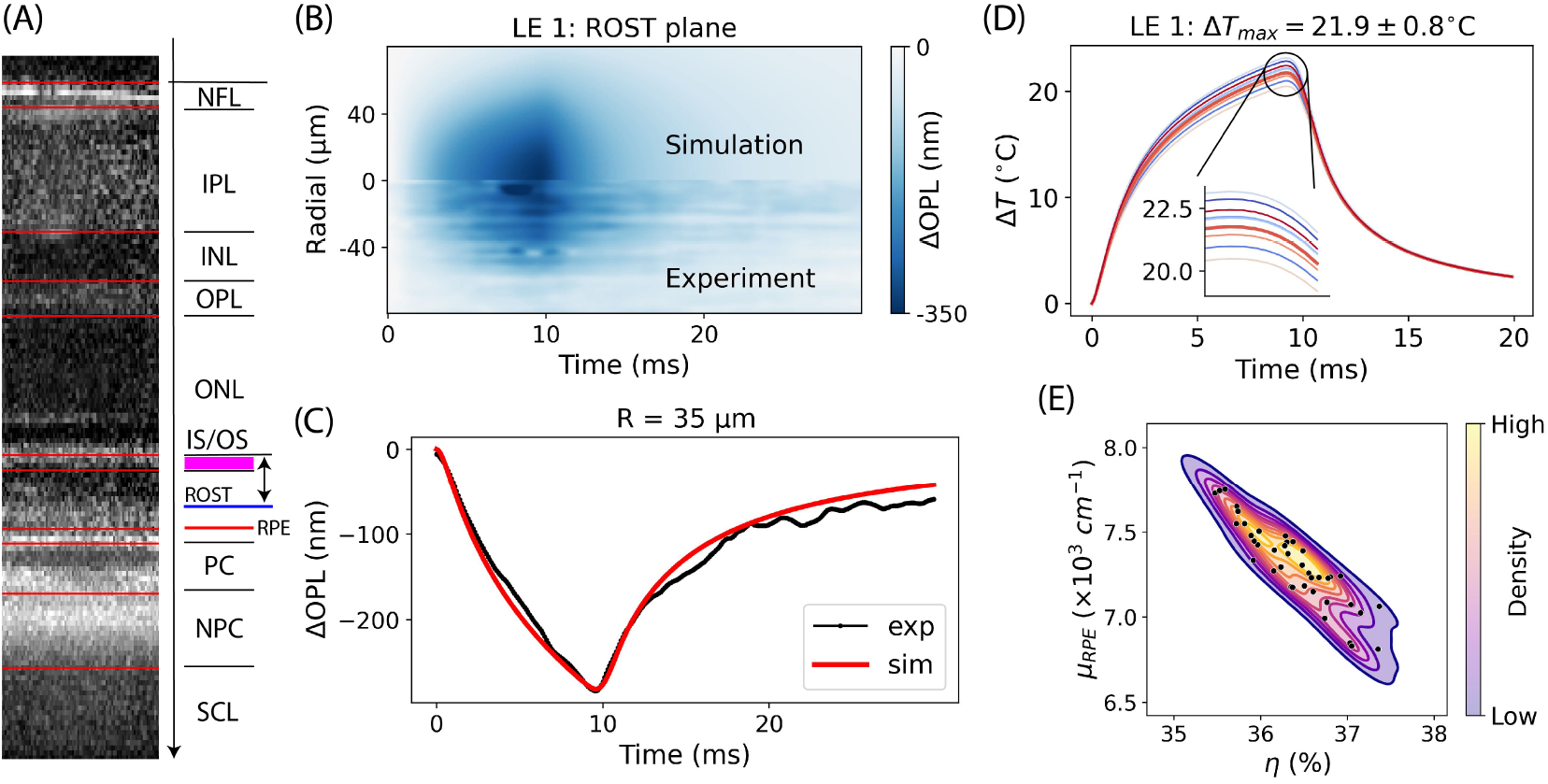
Comparison between the model and experimental data. (A) WT retinal layer segmentation and denotation. Distance between the center of the IS/OS line and the blue plane where the OPL change was extracted was set the same in simulation and in experiment. (B) Spatio-temporal lateral-referenced ΔOPL map compared between the best-fit simulation (top) and experiment (bottom). (C) An example trace of ΔOPL showing the heating and cooling dynamics. (D) Temperature time courses at the RPE computed with parameters collected from the last 10 iterations of the optimization process. (E) Contour plot showing the correlation between the absorption coefficient and optical transmittance via the bootstrapping analysis.

For parameter fitting optimization, the Nelder-Mead algorithm was utilized (45), which sometimes overshoots the parameters to avoid trapping in a local minimum. To ensure robustness of the solution, 3 or 4 different initial points were tested, and the one which converges to the lowest objective function value was selected. The last 10 iterations of the optimization process near the minima were then used to compute parameter uncertainties and the peak temperature variations at the RPE layer (Fig. 3D). The derived temperature has uncertainty less than 1 ^*°*^C, as listed in Table 1. Naturally, the optical transmittance and the absorption coefficient (equation 1) are strongly coupled, with the product of the two (i.e., *η × C*_*µ*_) representing the absorbed energy.

Using the model-predicted temperature profiles (Fig. 3D), it is possible to compute the associated Arrhenius integral and its uncertainty (Table 1). Heating of LE 1 retina was within the therapeutic window (0.1 *<* Ω *<* 1), whereas the heating of LE 2-3 was performed at lower input power and resulted in sub-therapeutic Arrhenius integral Ω *<* 0.1.

Temperature uncertainties can also be evaluated using a bootstrapping method (46, 47), and the animal LE 1 was randomly selected for this analysis. By adding 50 sets of noise to the best-fit model-predicted ΔOPL data, the model was then refit to these noisy data to obtain the new best-fit parameters and associated peak temperature (see *Supplementary Information*). The 50 tuples of (*η, C*_*µ*_)_*\**_ obtained from fitting the bootstrapping data shows a highly eccentric elliptical shape in the contour plot (Fig. 3E) confirming that these parameters are strongly and negatively correlated. The resulting temperature courses had lower standard deviation of the peak temperature (see Fig. S5C) than values listed in Table 1.

## Conclusions

Measurements of the phase changes in OCT induced by subtherapeutic tissue heating enables precise and safe temperature calibration for retinal laser therapy. In the future, when OCT machines will be upgraded to phase-sensitive detection, such mapping could be performed by the same tool as the structural OCT, and laser settings could be adjusted automatically during the treatment according to these maps using computer-guided scanning laser systems. Highly confined OPL change in WT retina, unlike the degenerate retina, suggests that cilium connecting the IS and OS in photoreceptors absorbs mechanical deformation induced by the thermal expansion. This feature might have diagnostic value.

## Materials and Methods

### Phase-sensitive OCT imaging

The high-speed line-scan spectral-domain OCT was assembled according to the optical layout shown in Fig. 4. The output of the supercontinuum laser (NKT FIR-9) was collimated (CL1, AC254-060-B) and filtered by two spectral filters (Semrock: FF01-776/LP-25, Thorlabs: FESH0900) yielding a bandwidth with a full-width at half-maximum (FWHM) of about 120 nm, centered at 840 nm.

**Fig. 4.**
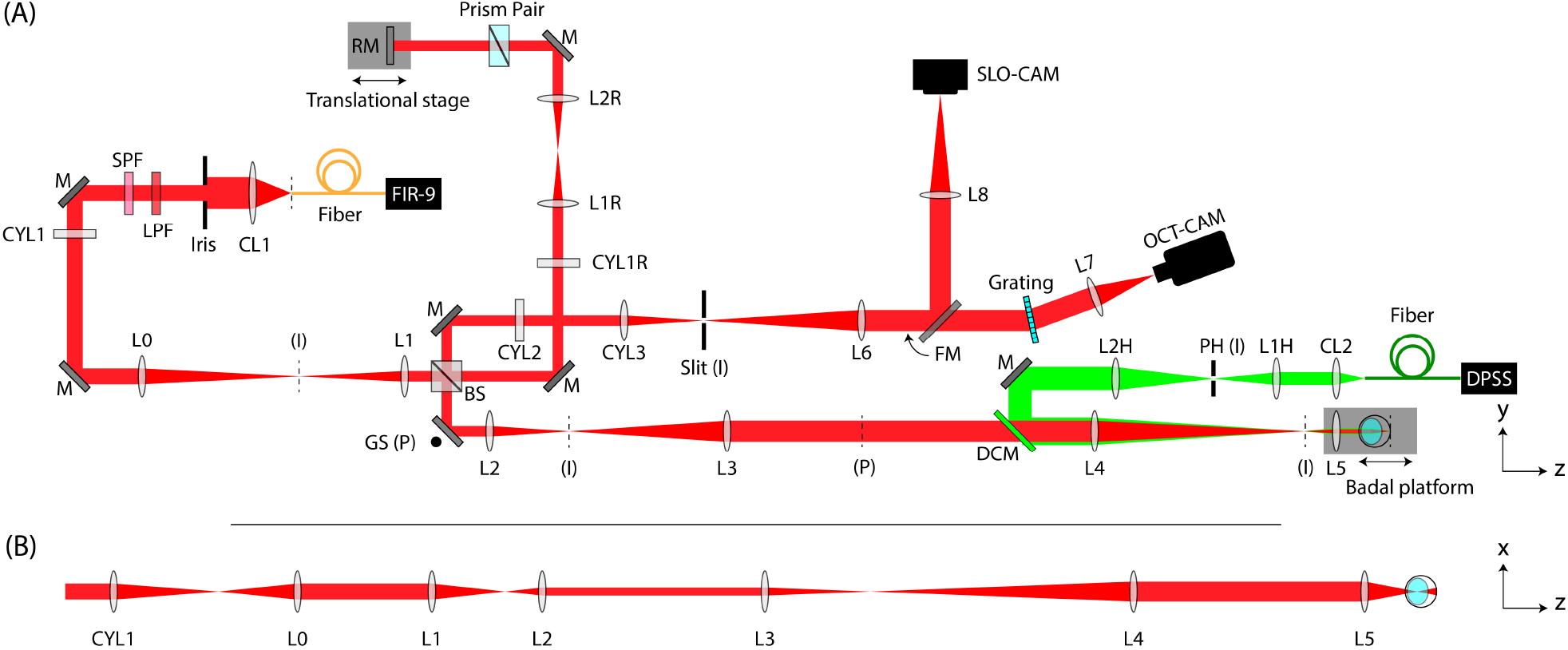
Phase-sensitive OCT setup. (A) Top view (*y*-*z*plane) including the line-scan spectral-domain OCT (red path) and the heating laser (green path). CL: collimating lens. SPF: short pass filter. LPF: long pass filter. CYL: achromatic cylindrical lens doublet. L: achromatic lens doublet. M: mirror. RM: reference mirror. BS: beamsplitter. GS: galvo scanner. DCM: dichroic mirror. FM: flippable mirror. PH: pinhole. DPSS: diode-pumped solid-state laser. FIR-9: NKT FIR-9 OCT laser. The effective focal lengths: CL1=60 mm, CYL1=CYL1R=CYL2=L4=L8=250 mm, L0=150 mm, L1=L1H=L7=100 mm, L2=CYL3=75 mm, L3=L1R=L2R=L2H=L6=200 mm, CL2=L5=30 mm. (I): conjugate image planes. (P): conjugate pupil planes. Not to scale. (B) Unfolded *x*-*z* view of the OCT illumination path showing line-field on the final retinal plane.

A cylindrical lens (CYL1) focuses the beam along the *x*-axis forming a line along the *y*-axis at its focal plane. After optical conjugation of the afocal telescope L0-L1, the beam was split into reference and sample paths using a 30:70 (R:T) nonpolarizing beamsplitter (Thorlabs, BSS11). In the sample arm, a one-dimensional galvo scanning (GS) mirror (8310K Series Galvanometer, Cambridge Technology), which steers the beam in the *y*-direction, was aligned at the system pupil plane followed by two afocal telescopes, L2-L3 and L4-L5. The lens L5 and the animal stage were placed together on the Badal platform to adjust beam focus on the retina. In the reference arm, a cylindrical lens (CYL1R) re-collimated the elliptical beam into a circular beam followed by an afocal telescope (L1R-L2R). A prism pair (#43-649, Littrow Dispersion Prism, Edmund Optics) was used to empirically balance the dispersion mismatch between the two arms during the experiment. For the detection path, the image plane was anamorphically conjugated to the slit using two cylindrical lenses (CYL2 and CYL3) and L2. The back-scattered light was diffracted by a 600 l/mm grating (WP-600/840-35×45, Wasatch Photonics) and finally focused on to the detector of the high-speed camera (Phantom v641) with an image size of 768*×*512 pixels (spectral*×*spatial). Note that the scanning laser ophthalmoscope (SLO) camera was aligned to be parfocal with the OCT camera and it was used to find a focal plane in the retina using Badal stage before the pOCT imaging.

The heating beam path was coupled with the OCT illumination path using a dichroic mirror (DCM, NFD01-532-25*×*36, Semrock) and was aligned to be coaxial with the OCT illumination. The heating beam source was a 532-nm diode-pumped solid-state laser (DPSS, 85-GHS-305-042, Melles Griot) coupled into an optical fiber (NA=0.22, diameter = 400 µm). The output of the fiber was collimated by CL2 and the tip of the fiber was conjugated to a pinhole with a diameter of 500-µm. The pinhole was demagnified and conjugated to OCT system image planes via the afocal telescope L2H-L4.

### Animal preparation

All experimental procedures were approved by the Stanford Administrative Panel on Laboratory Animal Care and conducted in accordance with the institutional guidelines and conformed to the Statement for the Use of Animals in Ophthalmic and Vision research of the Association for Research in Vision and Ophthalmology (ARVO). LE rats (P60 - P180) were used as a model of healthy retina and RCS rats were used as a model of outer retinal degeneration (22). All RCS rats were aged until P120 to allow for complete photoreceptor degeneration, as validated by OCT (HRA2-Spectralis; Heidelberg Engineering, Heidelberg, Germany). Animal colonies were maintained at the Stanford Animal Facility in 12-hour light/dark cycles with food and water ad libitum. Animals were anesthetized with a mixture of ketamine (75 mg/kg) and xylazine (5 mg/kg) injected intraperitoneally. The pupils were dilated with a mixture of 2.5% Phenylephrine Hydrocloride and 0.5% Tropicamide (Bausch & Lomb, Rochester, NY) ophthalmic solution and a zero power contact lens (base curvature 3.00 mm, diameter 6.00 mm, optical power 0.00 D; Lakewood, CO 80226) was placed on the eye for imaging. Animals were placed on a custom animal stage and secured with a bite-bar and ear-bars to minimize movement.

### Experimental ΔOPL data processing

We reconstructed complex-value OCT images from raw interferometric signals using *k*-linearity and discrete Fourier transform. To minimize decorrelation noise between repeated B-scans, we corrected bulk tissue motion with subpixel precision. Specifically, we estimated subpixel-level bulk displacements between first and the subsequent B-scans by locating the peak of upsampled cross-correlation maps (48). Each B-scan was then registered to the first B-scan using the phase-restoring subpixel image registration algorithm, which shifts the complex-value OCT image over arbitrary displacements (28). The registered B-scans were subsequently flattened along the BrM.

To further enhance phase stability, we self-referenced the phase changes to a region least affected by the stimulus the edges of the B-scan and extracted the OPL change in tissue following the laser pulse relative to the edges (14, 20). To cancel out the arbitrary phase offset at each pixel, we first computed the multiplication of each B-scan with the complex conjugate of the first B-scan

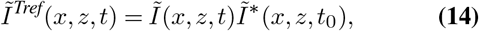

where *Ĩ* (*x, z, t*) is the complex-valued OCT signal after image registration and flattening. *Ĩ*_*Tref*_(*x, z, t*) is the corresponding time referenced signal where *x, z*, and *t* denote the indices along the lateral, axial and temporal dimensions, respectively. *t*_0_ denotes the time of the first frame, *\** represents complex conjugate. The time referenced signal was further averaged across certain depth range to enhance the signal-to-noise ratio (SNR). In this study, we averaged the complex-valued signal from NFL to the BrM and then extracted the phase component

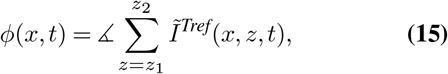

where *z*_1_ and *z*_2_ are the axial indices of the upper boundary of NFL and the lower boundary BrM, respectively. Note that *ϕ*(*x, t*) consisted of both phase changes following laser heating and residual phase fluctuations due to bulk tissue motion. In ex vivo studies, a single reference point far from the heating beam was commonly selected to remove undesired phase fluctuations (14, 19, 20). However, this strategy cannot be directly applied in vivo, as the rotational movement of the tissue resulted in varying phase along the lateral direction. Assuming the bulk motion, including both rotational and translational components, was rigid, the phase fluctuations should be linear along the lateral dimension (17). Accordingly, phase signals *ϕ*(*x*_*L*_, *t*) from a sequence of lateral positions *x*_*L*_ at the boundary of the OCT image (50 pixels on each side), which were least affected by the heating process, were selected to characterize the lateral phase shift. Specifically, *ϕ*(*x, t*) was first unwrapped along the lateral direction, resulting in *ϕ*^*u*^(*x, t*). We then conducted a linear fitting between *ϕ*^*u*^(*x*_*L*_, *t*) and *x*_*L*_ to extract phase shift *ϕ*^*fit*^(*x, t*) = *p*_0_(*t*)*x* + *p*_1_(*t*), where *p*_0_(*t*) and *p*_1_(*t*) are coefficients of the linear fitting. Hence, the lateral-referenced OCT signal can be found as

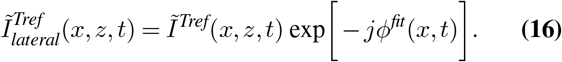

where *j* is the imaginary unit. We applied a Gaussian filter (kernel size: 5 × 7 pixels) on the complex-valued signal to enhance the SNR and then extract lateral-referenced phase signals,

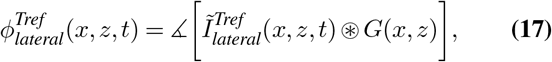

where *G* is the Gaussian kernel and ⊛ denotes the convolution operation. Finally, the laterally referenced ΔOPL can be computed as

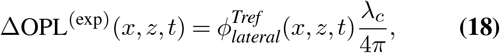

where *λ*_*c*_ = 840 nm denotes the central wavelength of the OCT imaging beam.

### Model-derived ΔOPL computation

For the axisymmetric model, we denote the lateral dimension as *r*. The variables of interest for ΔOPL computation from the model are *u*_*z*_(*r, z, t*) and Θ(*r, z, t*). For simplicity, we assume that the refractive index of the retinal tissue follows the same temperature dependence as of water. Hence, the single-pass OPL change at any point in space and time can be approximated as

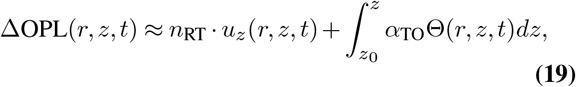

whe re *n*_RT_ is the refractive index of tissue at room temperature, and *α*_TO_ is the thermo-optic coefficient. Within the temperature range of interest, 37 *∼* 60*°*C, the temperature dependence of *n*_water_(*T*) varies between *−*1.4 *×* 10_*−*4_ and *−*1.8 *×* 10_*−*4_ K^*−*1^ (43). For simplicity, a linear variation with a slop of *α*_TO_ = *−*1.6 *×* 10^*−*4^ K^*−*1^ was used in this work. Refractive index at ambient temperature was taken to be 1.37 (49). As in the experimental data processing, the model-derived OPL changes were laterally referenced to the boundaries of the OCT B-scan.

## ACKNOWLEDGEMENTS

We thank Prof. Robert Zawadzki and Dr. Bingyao Tan for fruitful discussions of the optical setup. This work was funded by the National Institutes of Health (U01 EY032055), Air Force Office of Scientific Research (FA9550-20-1-0186) and Research to Prevent Blindness.

