## Supplementary Information for "Retinal thermometry in-vivo using phase-sensitive optical coherence tomography"

#### This PDF file includes:

Supporting text
Figs. S1 to S5
Table S1
Legends for Movies S1 to S2
SI References

#### Other supporting materials for this manuscript include the following:

Movies S1 to S2

### Supporting Information Text

**The Hankel-Laplace transform** The $m^{\text{th}}$-order Hankel-Laplace (HL) transform of a function $f$ from the primal domain $(r, z, t)$ to the HL domain $(\xi, z, s)$ is defined as (1):

$$\tilde{f}^{(m)}(\xi, z, s) = \int_0^\infty \int_0^\infty f(r, z, t)\, J_m(\xi r)\, e^{-st} r\, dr\, dt. \quad [1]$$

where $r$, $z$, $t$ denote the radial, axial, and the temporal dimensions, respectively. The associated $m^{\text{th}}$-order inverse HL transform is defined as

$$f(r, z, t) = \frac{1}{2\pi i} \int_{c-i\infty}^{c+i\infty} \int_0^\infty \tilde{f}^{(m)}(\xi, z, s)\, J_m(\xi r)\, e^{st}\, \xi\, d\xi\, ds. \quad [2]$$

Note three important identities of the Hankel transform ($\mathscr{H}_m$) used in the derivation of the thermo-mechanical retina model:

$$\frac{1}{r}\frac{\partial}{\partial r}\left(r\frac{\partial f}{\partial r}\right) - \frac{m^2}{r^2} f \xleftrightarrow{\mathscr{H}_m} -\xi^2 \tilde{f}^{(m)}, \quad [3]$$

$$r^{m-1}\frac{d}{dr}\left(r^{1-m} f\right) \xleftrightarrow{\mathscr{H}_m} -\xi \tilde{f}^{(m-1)}, \quad [4]$$

$$r^{-1-m}\frac{d}{dr}\left(r^{1+m} f\right) \xleftrightarrow{\mathscr{H}_m} \xi \tilde{f}^{(m+1)}. \quad [5]$$

Setting $m = 1$ and $m = 0$ for the last two identities, respectively,

$$\frac{df}{dr} \xleftrightarrow{\mathscr{H}_1} -\xi \tilde{f}^{(0)}, \quad [6]$$

$$\frac{1}{r}\frac{d}{dr}(rf) \xleftrightarrow{\mathscr{H}_0} \xi \tilde{f}^{(1)}. \quad [7]$$

### Derivation of the thermo-mechanical retina model

***The thermo-mechanical problem*** The rodent retina is modeled as an axisymmetric multi-layered medium. The physics of each layer is governed by thermo-elastic partial differential equations (PDE). In cylindrical coordinates, the equations of motion for an elastic medium in quasi-static conditions are (2):

$$\frac{\partial \sigma_r}{\partial r} + \frac{\partial \tau_{rz}}{\partial z} + \frac{\sigma_r - \sigma_\theta}{r} = 0, \quad [8]$$

$$\frac{\partial \tau_{rz}}{\partial r} + \frac{\partial \sigma_z}{\partial z} + \frac{\tau_{rz}}{r} = 0, \quad [9]$$

where $\sigma_r$, $\sigma_\theta$, and $\sigma_z$ are the normal stress components, $\tau_{rz}$ is the shear stress in the $r - z$ plane. The isotropic stress-strain constitutive relations (the generalized Hooke's law) including the thermal stress are:

$$\sigma_r + \beta\Theta = \lambda\epsilon_v + 2G\frac{\partial u_r}{\partial r}, \quad [10]$$

$$\sigma_\theta + \beta\Theta = \lambda\epsilon_v + 2G\frac{u_r}{r}, \quad [11]$$

$$\sigma_z + \beta\Theta = \lambda\epsilon_v + 2G\frac{\partial u_z}{\partial z}, \quad [12]$$

$$\tau_{rz} = G\left(\frac{\partial u_r}{\partial z} + \frac{\partial u_z}{\partial r}\right), \quad [13]$$

where

- $G$ is the shear modulus:

$$G = \frac{E}{2(1+\nu)} \quad [14]$$

  with $E$ the Young's modulus and $\nu$ the Poisson's ratio.

- $\lambda$ is the Lamé's first parameter:

$$\lambda = \frac{2G\nu}{1-2\nu}. \quad [15]$$

- $u_r$ and $u_z$ denote radial and axial displacements, respectively.

- $\epsilon_v$ is the volumetric strain:

$$\epsilon_v = \frac{\partial u_r}{\partial r} + \frac{u_r}{r} + \frac{\partial u_z}{\partial z}. \quad [16]$$

- $\beta\Theta$ is the thermal stress, where $\beta$ is the thermo-mechanical coupling parameter

$$\beta = 2G\alpha_{\text{TE}}\frac{1+\nu}{1-2\nu}, \quad [17]$$

  and $\Theta = T - T_0$ denotes the temperature change. Note $\alpha_{\text{TE}}$ is the thermal expansion coefficient.

The relationship between displacements and temperature change can be derived by substituting $\sigma_r$, $\sigma_\theta$, $\sigma_z$ and $\tau_{rz}$ from constitutive equations (Eqs. [10] to [13]) into the equations of motion (Eqs. [8] and [9]):

$$\nabla^2 u_r + \frac{1}{1-2\nu}\frac{\partial \epsilon_v}{\partial r} - \frac{1}{r^2}u_r - \frac{\beta}{G}\frac{\partial \Theta}{\partial r} = 0, \quad [18]$$

$$\nabla^2 u_z + \frac{1}{1-2\nu}\frac{\partial \epsilon_v}{\partial z} - \frac{\beta}{G}\frac{\partial \Theta}{\partial z} = 0, \quad [19]$$

where $\nabla^2$ is the Laplacian operator. The temperature change is governed by the Fourier heat conduction law and the heat diffusion equation:

$$\mathbf{q} = -\kappa\nabla\Theta, \quad [20]$$

$$\frac{\partial \Theta}{\partial t} = \alpha\nabla^2\Theta, \quad [21]$$

where $\mathbf{q} = [q_r, q_z]^\intercal$ is the heat flux vector, $\kappa$ is the coefficient of thermal conductivity, $\alpha = \kappa/\rho c_p$ is the coefficient of thermal diffusivity, $\rho$ is the material density, $c_p$ is the material-specific heat capacity, and $\nabla$ is the gradient operator.

***The state matrix equation*** With equations [18], [19] and [21], the state equations of an elastic layer can be derived. Taking the 1st-order HL transform of equation [18] gives:

$$\frac{\partial^2 \tilde{u}_r^{(1)}}{\partial z^2} - \frac{2(1-\nu)}{1-2\nu}\tilde{u}_r^{(1)} - \frac{1}{1-2\nu}\xi\frac{\partial \tilde{u}_z^{(0)}}{\partial z} + \frac{\beta}{G}\xi\tilde{\Theta}^{(0)} = 0, \tag{22}$$

where HL property [6] has been used to convert the axial displacement variable and temperature change variable in the HL domain from 1st-order to 0th-order. Similarly, taking the 0th-order HL transform of equation [19] gives:

$$\frac{2(1-\nu)}{1-2\nu}\frac{\partial^2 \tilde{u}_z^{(0)}}{\partial z^2} - \xi^2\tilde{u}_z^{(0)} + \frac{1}{1-2\nu}\xi\frac{\partial \tilde{u}_r^{(1)}}{\partial z} - \frac{\beta}{G}\frac{\partial \tilde{\Theta}^{(0)}}{\partial z} = 0. \tag{23}$$

Lastly, taking the 0th-order HL transform of equation [21] gives:

$$(s+\alpha\xi^2)\tilde{\Theta}^{(0)} - \alpha\frac{\partial^2 \tilde{\Theta}^{(0)}}{\partial z^2} = 0. \tag{24}$$

Define $\mathbf{\Lambda}$ as the state vector:

$$\mathbf{\Lambda} = \begin{bmatrix} \tilde{u}_r^{(1)} \\ \tilde{u}_z^{(0)} \\ \tilde{\Theta}^{(0)} \end{bmatrix} \tag{25}$$

and let $\mathbf{W} = \begin{bmatrix} \mathbf{\Lambda} \\ \partial\mathbf{\Lambda}/\partial z \end{bmatrix}$, three state equations can be combined into one matrix equation:

$$\frac{\partial \mathbf{W}(\xi,z,s)}{\partial z} = \mathbf{\Phi}(\xi,s)\mathbf{W}(\xi,z,s). \tag{26}$$

where $\mathbf{\Phi} \in \mathbb{R}^{6\times 6}$ is given below:

$$\mathbf{\Phi}(\xi,s) = \begin{bmatrix} 0 & 0 & 0 & 1 & 0 & 0 \\ 0 & 0 & 0 & 0 & 1 & 0 \\ 0 & 0 & 0 & 0 & 0 & 1 \\ \frac{2(1-\nu)}{1-2\nu}\xi^2 & 0 & -\frac{\beta}{G}\xi & 0 & \frac{1}{1-2\nu}\xi & 0 \\ 0 & \frac{1-2\nu}{2(1-\nu)}\xi^2 & 0 & -\frac{1}{2(1-\nu)}\xi & 0 & \frac{1-2\nu}{2(1-\nu)}\frac{\beta}{G} \\ 0 & 0 & \frac{s}{\alpha}+\xi^2 & 0 & 0 & 0 \end{bmatrix} \tag{27}$$

The eigenvalues of $\mathbf{\Phi}$ are found as

$$\lambda_{1,2} = \xi, \tag{28}$$

$$\lambda_{3,4} = -\xi, \tag{29}$$

$$\lambda_5 = \sqrt{\frac{s}{\alpha}+\xi^2}, \tag{30}$$

$$\lambda_6 = -\sqrt{\frac{s}{\alpha}+\xi^2}. \tag{31}$$

Corresponding eigenvectors are found as columns of the following matrix:

$$\begin{bmatrix} -\xi^{-1} & -2(1-2\nu)\xi^{-2} & -\xi^{-1} & 2(1-2\nu)\xi^{-2} & \eta\cdot p^{-1} & -\eta\cdot p^{-1} \\ \xi^{-1} & -\xi^{-2} & -\xi^{-1} & -\xi^{-2} & \eta & \eta \\ 0 & 0 & 0 & 0 & -p^{-1} & p^{-1} \\ -1 & (-3+4\nu)\xi^{-1} & 1 & (-3+4\nu)\xi^{-1} & -\eta\cdot\xi & -\eta\cdot\xi \\ 1 & 0 & 1 & 0 & -\eta\cdot p & \eta\cdot p \\ 0 & 0 & 0 & 0 & 1 & 1 \end{bmatrix}, \tag{32}$$

where

$$\eta = \frac{(1-2\nu)\alpha\beta}{2(1-\nu)sG}, \tag{33}$$

$$p = \sqrt{\frac{s}{\alpha}+\xi^2}. \tag{34}$$

Note that the eigenvectors in [32] are generalized eigenvectors solved using Jordan decomposition. Denote the $i^{th}$ generalized eigenvectors as $\mathbf{m}_i$, the general solution to the first-order matrix differential state equation [26] with algebraic multiplicity in eigenvalues has the form of

$$\mathbf{W}(\xi,z,s) = \mathbf{m}_1 e^{\lambda_1 z}\cdot\ell_1 + (\mathbf{m}_1 z+\mathbf{m}_2)e^{\lambda_2 z}\cdot\ell_2 + \mathbf{m}_3 e^{\lambda_3 z}\cdot\ell_3 + (\mathbf{m}_3 z+\mathbf{m}_4)e^{\lambda_4 z}\cdot\ell_4 + \mathbf{m}_5 e^{\lambda_5 z}\cdot\ell_5 + \mathbf{m}_6 e^{\lambda_6 z}\cdot\ell_6 \quad [35]$$

$$= \begin{bmatrix}\mathbf{m}_1 e^{\xi z} & (\mathbf{m}_1 z+\mathbf{m}_2)e^{\xi z} & \mathbf{m}_3 e^{-\xi z} & (\mathbf{m}_3 z+\mathbf{m}_4)e^{-\xi z} & \mathbf{m}_5 e^{\sqrt{\frac{s}{a}+\xi^2}z} & \mathbf{m}_6 e^{-\sqrt{\frac{s}{a}+\xi^2}z}\end{bmatrix}\begin{bmatrix}\ell_1\\ \ell_2\\ \ell_3\\ \ell_4\\ \ell_5\\ \ell_6\end{bmatrix} \quad [36]$$

$$= \tilde{\mathbf{M}}(\xi,z,s)\mathbf{L}, \quad [37]$$

where $\ell_i$'s are arbitrary constants. Consider a single finite layer with the top boundary at depth $z=0$ and the bottom boundary at depth $z$, and take the top half of the vector $\mathbf{W}$ and the matrix $\tilde{\mathbf{M}}$, the states at the boundaries can be expressed as

$$\begin{bmatrix}\mathbf{\Lambda}(\xi,0,s)\\ \mathbf{\Lambda}(\xi,z,s)\end{bmatrix} = \begin{bmatrix}\mathbf{M}(\xi,0,s)\\ \mathbf{M}(\xi,z,s)\end{bmatrix}\mathbf{L}, \quad [38]$$

where $\mathbf{M}$ is the top half of the matrix $\tilde{\mathbf{M}}$, i.e.,

$$\mathbf{M}(\xi,z,s) = \begin{bmatrix} -\xi^{-1}e^{\xi z} & -(\xi^{-1}z+2(1-2\nu)\xi^{-2})e^{\xi z} & -\xi^{-1}e^{-\xi z} & (-\xi^{-1}z+2(1-2\nu)\xi^{-2})e^{-\xi z} & \eta\cdot p^{-1}e^{pz} & -\eta\cdot p^{-1}e^{-pz}\\ \xi^{-1}e^{\xi z} & (\xi^{-1}z-\xi^{-2})e^{\xi z} & -\xi^{-1}e^{-\xi z} & -(\xi^{-1}z+\xi^{-2})e^{-\xi z} & \eta\, e^{pz} & \eta\, e^{-pz}\\ 0 & 0 & 0 & 0 & -p^{-1}e^{pz} & p^{-1}e^{-pz}\end{bmatrix} \quad [39]$$

***The stress matrix equation*** The normal stress along $z-$axis, the shear stress in the $r-z$ plane, and the heat flow are chosen to form the stress vector in HL domain, i.e.,

$$\mathbf{V} = \begin{bmatrix}\tilde{\tau}_{rz}^{(1)}\\ \tilde{\sigma}_z^{(0)}\\ \tilde{Q}^{(0)}\end{bmatrix} \quad [40]$$

The relationship between the stress vector and the state vector can be derived from equations [12-13] and the heat flow in the $z-$direction based on the Fourier heat conduction law:

$$Q = \int_0^t q_z\, d\tau = -\int_0^t \kappa\frac{\partial\Theta}{\partial z}\, d\tau. \quad [41]$$

Taking the HL transform of equations [12-13] and [41] gives

$$\tilde{\tau}_{rz}^{(1)} = G\left(\frac{\partial \tilde{u}_r^{(1)}}{\partial z} - \xi\tilde{u}_z^{(0)}\right), \quad [42]$$

$$\tilde{\sigma}_z^{(0)} = \frac{2G\nu}{1-2\nu}\xi\tilde{u}_r^{(1)} + 2G\frac{1-\nu}{1-2\nu}\frac{\partial\tilde{u}_z^{(0)}}{\partial z} - \beta\tilde{\Theta}^{(0)}, \quad [43]$$

$$\tilde{Q}^{(0)} = -\frac{\kappa}{s}\frac{\partial\tilde{\Theta}^{(0)}}{\partial z}. \quad [44]$$

Hence, the stress matrix equation is derived:

$$\mathbf{V}(\xi,z,s) = \mathbf{S}(\xi,s)\mathbf{W}(\xi,z,s), \quad [45]$$

where $\mathbf{S}\in\mathbb{R}^{3\times 6}$:

$$\mathbf{S}(\xi,s) = \begin{bmatrix} 0 & -G\xi & 0 & G & 0 & 0\\ \frac{2G\nu}{1-2\nu}\xi & 0 & -\beta & 0 & 2G\frac{1-\nu}{1-2\nu} & 0\\ 0 & 0 & 0 & 0 & 0 & -\frac{\kappa}{s}\end{bmatrix} \quad [46]$$

As derived in the previous section, $\mathbf{W}(\xi,z,s) = \tilde{\mathbf{M}}(\xi,z,s)\mathbf{L}$, hence

$$\mathbf{V}(\xi,z,s) = \mathbf{S}(\xi,s)\tilde{\mathbf{M}}(\xi,z,s)\mathbf{L}. \quad [47]$$

Denote $\mathbf{N}(\xi,z,s) = \mathbf{S}(\xi,s)\tilde{\mathbf{M}}(\xi,z,s)$, for a single finite layer with the top boundary at depth $z=0$ and the bottom boundary at depth $z$, we have

$$\begin{bmatrix}-\mathbf{V}(\xi,0,s)\\ \mathbf{V}(\xi,z,s)\end{bmatrix} = \begin{bmatrix}-\mathbf{N}(\xi,0,s)\\ \mathbf{N}(\xi,z,s)\end{bmatrix}\mathbf{L}. \quad [48]$$

The reason for using the minus sign will become clear later.

***The stiffness matrix of a single elastic layer*** Combining equations [38] and [48] to eliminate $\mathbf{L}$ yields

$$\begin{bmatrix} -\mathbf{V}(\xi,0,s) \\ \mathbf{V}(\xi,z,s) \end{bmatrix} = \mathbf{K}(\xi,z,s) \begin{bmatrix} \mathbf{\Lambda}(\xi,0,s) \\ \mathbf{\Lambda}(\xi,z,s) \end{bmatrix}, \tag*{[49]}$$

where matrix $\mathbf{K} \in \mathbb{R}^{6\times 6}$ is called the stiffness matrix

$$\mathbf{K}(\xi,z,s) = \begin{bmatrix} -\mathbf{N}(\xi,0,s) \\ \mathbf{N}(\xi,z,s) \end{bmatrix} \begin{bmatrix} \mathbf{M}(\xi,0,s) \\ \mathbf{M}(\xi,z,s) \end{bmatrix}^{-1}. \tag*{[50]}$$

Equation [49] is referred to as the transfer matrix equation in this modeling. Elements in $\mathbf{K}$, also given in (3), are:

$$\mathbf{K}_{11} = \mathbf{K}_{44} = \frac{2G\xi b_2(4z\xi e_1^2 b_1 - b_3 d_1 d_3)}{g}, \tag*{[51]}$$

$$\mathbf{K}_{12} = \mathbf{K}_{21} = -\mathbf{K}_{45} = -\mathbf{K}_{54} = \frac{2G\xi(4z^2\xi^2 e_1^2 b_1^2 - Gb_3 d_1^2)}{g}, \tag*{[52]}$$

$$\mathbf{K}_{13} = \mathbf{K}_{46} = -\frac{2G\beta\alpha\xi[4z\xi e_1 b_1(p\, e_2 d_1 - \xi e_1 d_2) + b_3 d_1(\xi d_2 d_3 - p\, d_1 d_4)]}{s\, g\, d_2}, \tag*{[53]}$$

$$\mathbf{K}_{14} = \mathbf{K}_{41} = \frac{4G\xi e_1 b_2(b_3 d_1 - z\xi b_1 d_3)}{g}, \tag*{[54]}$$

$$\mathbf{K}_{15} = \mathbf{K}_{51} = -\mathbf{K}_{24} = -\mathbf{K}_{42} = -\frac{4Gz\xi^2 e_1 b_1 b_2 d_1}{g}, \tag*{[55]}$$

$$\mathbf{K}_{16} = \mathbf{K}_{43} = -\frac{4G\beta\alpha\xi[z\xi e_1 b_1(\xi d_2 d_3 - p\, d_1 d_4) + b_3 d_1(p\, e_2 d_1 - \xi e_1 d_2)]}{s\, g\, d_2}, \tag*{[56]}$$

$$\mathbf{K}_{22} = \mathbf{K}_{55} = -\frac{2G\xi b_2(4z\xi e_1^2 b_1 + b_3 d_1 d_3)}{g}, \tag*{[57]}$$

$$\mathbf{K}_{23} = -\mathbf{K}_{56} = \frac{2G\beta\alpha\xi[4z\xi p\, e_1 b_1 f + 4p\, e_1 e_2 b_3 d_1 + b_3 d_1(\xi d_1 d_2 - p\, d_3 d_4)]}{s\, g\, d_2}, \tag*{[58]}$$

$$\mathbf{K}_{25} = \mathbf{K}_{52} = \frac{4G\xi e_1 b_2(b_3 d_1 + z\xi b_1 d_3)}{g}, \tag*{[59]}$$

$$\mathbf{K}_{26} = -\mathbf{K}_{53} = \frac{4G\beta\alpha\xi[4z\xi p\, e_1^2 e_2 b_1 + z\xi e_1 b_1(\xi d_1 d_2 - p\, d_3 d_4) + p\, b_3 d_1 f]}{s\, g\, d_2}, \tag*{[60]}$$

$$\mathbf{K}_{33} = \mathbf{K}_{66} = -\frac{\kappa p\, d_4}{s\, d_2}, \tag*{[61]}$$

$$\mathbf{K}_{36} = \mathbf{K}_{63} = \frac{2\kappa p\, e_2}{s\, d_2}, \tag*{[62]}$$

and $\mathbf{K}_{31} = \mathbf{K}_{32} = \mathbf{K}_{34} = \mathbf{K}_{35} = \mathbf{K}_{61} = \mathbf{K}_{62} = \mathbf{K}_{64} = \mathbf{K}_{65} = 0$, where

$$b_1 = \lambda + G \tag*{[63]}$$

$$b_2 = \lambda + 2G \tag*{[64]}$$

$$b_3 = \lambda + 3G \tag*{[65]}$$

$$e_1 = e^{-\xi z} \tag*{[66]}$$

$$e_2 = e^{-pz} \tag*{[67]}$$

$$d_1 = 1 - e_1^2 \tag*{[68]}$$

$$d_2 = 1 - e_2^2 \tag*{[69]}$$

$$d_3 = 1 + e_1^2 \tag*{[70]}$$

$$d_3 = 1 + e_2^2 \tag*{[71]}$$

$$f = e_2 d_3 - e_1 d_4 \tag*{[72]}$$

$$g = 4z^2\xi^2 e_1^2 b_1^2 - b_3^2 d_1^2. \tag*{[73]}$$

It is worth to write out the stiffness matrix and observe symmetry among its elements:

$$\mathbf{K}(\xi,z,s) = \begin{bmatrix} \mathbf{A}_{3\times3} & \mathbf{B}_{3\times3} \\ \mathbf{C}_{3\times3} & \mathbf{D}_{3\times3} \end{bmatrix} = \begin{bmatrix} \mathbf{K}_{11} & \mathbf{K}_{12} & \mathbf{K}_{13} & \mathbf{K}_{14} & \mathbf{K}_{15} & \mathbf{K}_{16} \\ \mathbf{K}_{12} & \mathbf{K}_{22} & \mathbf{K}_{23} & -\mathbf{K}_{15} & \mathbf{K}_{25} & \mathbf{K}_{26} \\ 0 & 0 & \mathbf{K}_{33} & 0 & 0 & \mathbf{K}_{36} \\ \mathbf{K}_{14} & -\mathbf{K}_{15} & \mathbf{K}_{16} & \mathbf{K}_{11} & -\mathbf{K}_{12} & \mathbf{K}_{13} \\ \mathbf{K}_{15} & \mathbf{K}_{25} & -\mathbf{K}_{26} & -\mathbf{K}_{12} & \mathbf{K}_{22} & -\mathbf{K}_{23} \\ 0 & 0 & \mathbf{K}_{36} & 0 & 0 & \mathbf{K}_{33} \end{bmatrix} \tag*{[74]}$$

Note the similarity between submatrices $\mathbf{A}$ and $\mathbf{D}$, also submatrices $\mathbf{B}$ and $\mathbf{C}$.

***The mathematical formulation of the retinal N-layer assembly*** The geometry and coordinate system of the $N$-layer assembly is defined in Fig. S1. The top plane of the retina is at $z = 0$, and the space above it is called the top half-space defined as the $1^{\text{st}}$ layer. Similarly, the last layer of the assembly will extend from $z = z_R$ ($z_R$:the thickness of the retina) to $z = +\infty$ and is called the bottom half-space. Denoting the stiffness matrix of the $i^{\text{th}}$ layer as

$$\mathbf{K}^{[i]} = \begin{bmatrix} \mathbf{A}^{[i]} & \mathbf{B}^{[i]} \\ \mathbf{C}^{[i]} & \mathbf{D}^{[i]} \end{bmatrix}, \tag{75}$$

then the transfer matrix equation for the $i^{\text{th}}$ layer can be written as two equations using the submatrices, i.e.,

$$-\mathbf{V}(\xi, z_i^+, s) = \mathbf{A}^{[i]}\mathbf{\Lambda}(\xi, z_i^+, s) + \mathbf{B}^{[i]}\mathbf{\Lambda}(\xi, z_{i+1}^-, s), \tag{76}$$

$$\mathbf{V}(\xi, z_{i+1}^-, s) = \mathbf{C}^{[i]}\mathbf{\Lambda}(\xi, z_i^+, s) + \mathbf{D}^{[i]}\mathbf{\Lambda}(\xi, z_{i+1}^-, s). \tag{77}$$

For the boundary conditions, we assume no stress, displacement, heat transfer, or temperature rise at infinity, i.e.,

$$\mathbf{V}(\xi, -\infty, s) = \mathbf{V}(\xi, +\infty, s) = \mathbf{\Lambda}(\xi, -\infty, s) = \mathbf{\Lambda}(\xi, +\infty, s) = \mathbf{0}_{1\times 3}. \tag{78}$$

With the boundary conditions, the transfer matrix equations for the top and bottom half-spaces are reduced to

$$\mathbf{V}(\xi, 0^-, s) = \mathbf{D}^{[1]}\mathbf{\Lambda}(\xi, 0^-, s), \tag{79}$$

$$-\mathbf{V}(\xi, z_R^+, s) = \mathbf{A}^{[N]}\mathbf{\Lambda}(\xi, z_R^+, s), \tag{80}$$

where

$$\mathbf{D}^{[1]} = \lim_{z\to-\infty} \begin{bmatrix} \mathbf{K}_{44}^{[1]} & \mathbf{K}_{45}^{[1]} & \mathbf{K}_{46}^{[1]} \\ \mathbf{K}_{54}^{[1]} & \mathbf{K}_{55}^{[1]} & \mathbf{K}_{56}^{[1]} \\ \mathbf{K}_{64}^{[1]} & \mathbf{K}_{65}^{[1]} & \mathbf{K}_{66}^{[1]} \end{bmatrix} = \begin{bmatrix} 0 & 0 & 0 \\ 0 & 0 & 0 \\ 0 & 0 & -\frac{\kappa p}{s} \end{bmatrix}, \tag{81}$$

$$\mathbf{A}^{[N]} = \lim_{z\to+\infty} \begin{bmatrix} \mathbf{K}_{11}^{[N]} & \mathbf{K}_{12}^{[N]} & \mathbf{K}_{13}^{[N]} \\ \mathbf{K}_{21}^{[N]} & \mathbf{K}_{22}^{[N]} & \mathbf{K}_{23}^{[N]} \\ \mathbf{K}_{31}^{[N]} & \mathbf{K}_{32}^{[N]} & \mathbf{K}_{33}^{[N]} \end{bmatrix} = \begin{bmatrix} 0 & 0 & 0 \\ 0 & 0 & 0 \\ 0 & 0 & -\frac{\kappa p}{s} \end{bmatrix}. \tag{82}$$

The continuity conditions at interfaces between layers without external forces or heat sources are given as:

$$\mathbf{V}(\xi, z_i^-, s) = \mathbf{V}(\xi, z_i^+, s) = \mathbf{V}(\xi, z_i, s), \tag{83}$$

$$\mathbf{\Lambda}(\xi, z_i^-, s) = \mathbf{\Lambda}(\xi, z_i^+, s) = \mathbf{\Lambda}(\xi, z_i, s). \tag{84}$$

At an interface where a surface heat source is present, e.g., at the surface of the RPE layer ($z = z_{\text{RPE}}$), the boundary condition is:

$$\mathbf{V}(\xi, z_{\text{RPE}}^+, s) = \mathbf{V}(\xi, z_{\text{RPE}}^-, s) + \mathbf{F}(\xi, z_{\text{RPE}}, s), \tag{85}$$

where $\mathbf{F}$ represents the external heat source term, i.e.,

$$\mathbf{F}(\xi, z_{\text{RPE}}, s) = \begin{bmatrix} 0 \\ 0 \\ \tilde{Q}_e^{(0)}(\xi, z_{\text{RPE}}, s) \end{bmatrix}, \tag{86}$$

where $\tilde{Q}_e^{(0)}(\xi, z_{\text{RPE}}, s)$ is the HL transform of the external heat flow $Q_e(r, z_{\text{RPE}}, t)$ brought through the laser absorption (see next section **Surface heat source**). Any layer has a system of matrix equations similar to [76] and [77], hence, for $N$ layers, there is a total of $2N - 2$ matrix equations (the top and bottom half-spaces only need 1 matrix equation for each). The introduction of the minus sign in equation [49] with the continuity conditions simplifies $2N - 2$ to $N - 1$ equations. For example, consider a simple 4-layer assembly, with the top half-space (layer 1, between $z_1 = -\infty$ and $z_2 = 0$), a water layer, an RPE layer, and the bottom half-space, with a surface heat source located at the top of the RPE layer, the complete set of matrix equations are

$$\mathbf{V}(\xi, 0^-, s) = \mathbf{D}^{[1]}\mathbf{\Lambda}(\xi, 0^-, s), \tag{87}$$

$$-\mathbf{V}(\xi, 0^+, s) = \mathbf{A}^{[2]}\mathbf{\Lambda}(\xi, 0^+, s) + \mathbf{B}^{[2]}\mathbf{\Lambda}(\xi, z_3^-, s), \tag{88}$$

$$\mathbf{V}(\xi, z_3^-, s) + \mathbf{F}(\xi, z_3, s) = \mathbf{C}^{[2]}\mathbf{\Lambda}(\xi, 0^+, s) + \mathbf{D}^{[2]}\mathbf{\Lambda}(\xi, z_3^-, s), \tag{89}$$

$$-\mathbf{V}(\xi, z_3^+, s) = \mathbf{A}^{[3]}\mathbf{\Lambda}(\xi, z_3^+, s) + \mathbf{B}^{[3]}\mathbf{\Lambda}(\xi, z_4^-, s), \tag{90}$$

$$\mathbf{V}(\xi, z_4^-, s) = \mathbf{C}^{[3]}\mathbf{\Lambda}(\xi, z_3^+, s) + \mathbf{D}^{[3]}\mathbf{\Lambda}(\xi, z_4^-, s), \tag{91}$$

$$-\mathbf{V}(\xi, z_4^+, s) = \mathbf{A}^{[4]}\mathbf{\Lambda}(\xi, z_4^+, s). \tag{92}$$

Combine [87-88], [89-90], and [91-92], and then apply continuity conditions to obtain three resultant equations:

$$\mathbf{0}_{3\times1} = (\mathbf{D}^{[1]} + \mathbf{A}^{[2]})\boldsymbol{\Lambda}(\xi, 0, s) + \mathbf{B}^{[2]}\boldsymbol{\Lambda}(\xi, z_3, s), \qquad [93]$$

$$\mathbf{F}(\xi, z_3, s) = \mathbf{C}^{[2]}\boldsymbol{\Lambda}(\xi, 0, s) + (\mathbf{D}^{[2]} + \mathbf{A}^{[3]})\boldsymbol{\Lambda}(\xi, z_3, s) + \mathbf{B}^{[3]}\boldsymbol{\Lambda}(\xi, z_4, s), \qquad [94]$$

$$\mathbf{0}_{3\times1} = \mathbf{C}^{[3]}\boldsymbol{\Lambda}(\xi, z_3, s) + (\mathbf{D}^{[3]} + \mathbf{A}^{[4]})\boldsymbol{\Lambda}(\xi, z_4, s), \qquad [95]$$

which can be written as one matrix equation:

$$\begin{bmatrix} \mathbf{0}_{3\times1} \\ \mathbf{F}(\xi, z_3, s) \\ \mathbf{0}_{3\times1} \end{bmatrix} = \begin{bmatrix} \mathbf{D}^{[1]} + \mathbf{A}^{[2]} & \mathbf{B}^{[2]} & \mathbf{0}_{3\times3} \\ \mathbf{C}^{[2]} & \mathbf{D}^{[2]} + \mathbf{A}^{[3]} & \mathbf{B}^{[3]} \\ \mathbf{0}_{3\times3} & \mathbf{C}^{[3]} & \mathbf{D}^{[3]} + \mathbf{A}^{[4]} \end{bmatrix} \begin{bmatrix} \boldsymbol{\Lambda}(\xi, 0, s) \\ \boldsymbol{\Lambda}(\xi, z_3, s) \\ \boldsymbol{\Lambda}(\xi, z_4, s) \end{bmatrix}. \qquad [96]$$

This simple example can be extended to a $N$-layer assembly. By inverting the final matrix equation, the state vector $\boldsymbol{\Lambda}(\xi, z_i, s)$ at any interface can be found. For brevity, the equation [96] is written as

$$\tilde{\mathbf{V}}(\xi, s) = \boldsymbol{\mathcal{K}}(\xi, s)\tilde{\boldsymbol{\Lambda}}(\xi, s). \qquad [97]$$

***Numerical inverse Hankel-Laplace transform*** To obtain $u_z(r, z, t)$ and $\Theta(r, z, t)$ for the $\Delta$OPL calculation from $\boldsymbol{\Lambda}(\xi, z, s)$, we perform numerical HL transform inversion of $\tilde{u}_z^{(0)}(\xi, z, s)$ and $\tilde{\Theta}^{(0)}(\xi, z, s)$. The inversion can be treated independently:

- The inverse Hankel transform is defined by the integral:

$$f(r) = \int_0^\infty \tilde{f}(\xi)J_m(\xi r)\xi d\xi, \qquad [98]$$

  which can be approximated as

$$f(r) = \sum_{i=0}^{\infty} \int_{\xi_i}^{\xi_{i+1}} \tilde{f}(\xi)J_m(\xi r)\xi d\xi, \qquad [99]$$

$$\approx \sum_{i=1}^{N_B} \int_{\xi_i}^{\xi_{i+1}} \tilde{f}(\xi)J_m(\xi r)\xi d\xi, \qquad [100]$$

  where $\xi_i$'s are zero points of the Bessel function $J_m$ and $N_B$ denotes the number of Bessel intervals considered. The integrals over the intervals $[\xi_i, \xi_{i+1}]$ can be approximated using the Gauss-Legendre quadrature:

$$\int_{\xi_i}^{\xi_{i+1}} \tilde{f}(\xi)J_m(\xi r)\xi d\xi \approx \frac{\xi_{i+1} - \xi_i}{2} \sum_{p=1}^{N_{GL}} w_p \tilde{f}(\beta_{i,p})J_m(r\beta_{i,p})\beta_{i,p}, \qquad [101]$$

  where $N_{GL}$ denotes the quadrature order, $w_p$ and $r_p$ are the Gaussian weights and nodes, respectively (4), and

$$\beta_{i,p} = \frac{\xi_{i+1} - \xi_i}{2} r_p + \frac{\xi_{i+1} + \xi_i}{2}. \qquad [102]$$

- The Abate-Whitt framework (5) is used for the inverse Laplace transform in which $f(t)$ is approximated as a linear combination of $\tilde{f}(s)$ values, i.e., for $t > 0$

$$f(t) \approx \sum_{k=1}^{N_L} \frac{\eta_k}{t} \tilde{f}\left(\frac{\gamma_k}{t}\right), \qquad [103]$$

  where nodes $\gamma_k$ and weights $\eta_k$ are $N_L$-dependent real or complex numbers. The nodes and weights are made available by the authors of (6) which uses concentrated matrix exponential distribution.

In summary, to obtain $u_z(r, z, t)$ and $\Theta(r, z, t)$, we need to compute

$$u_z(r, z, t) \approx \sum_{k=1}^{N_L} \frac{\eta_k}{t} \sum_{i=1}^{N_B} \frac{\xi_{i+1} - \xi_i}{2} \sum_{p=1}^{N_{GL}} w_p \tilde{u}_z^{(0)}\left(\beta_{i,p}, z, \frac{\gamma_k}{t}\right) J_m(r\beta_{i,p})\beta_{i,p}, \qquad [104]$$

and

$$\Theta(r, z, t) \approx \sum_{k=1}^{N_L} \frac{\eta_k}{t} \sum_{i=1}^{N_B} \frac{\xi_{i+1} - \xi_i}{2} \sum_{p=1}^{N_{GL}} w_p \tilde{\Theta}^{(0)}\left(\beta_{i,p}, z, \frac{\gamma_k}{t}\right) J_m(r\beta_{i,p})\beta_{i,p}, \qquad [105]$$

***The general algorithm*** The computational flow of equations [104-105] in MATLAB or Python is summarized in a pseudo-code format shown below:

**Algorithm 1** State inversion

```
Define arrays E, X, B, W, J, U_z, T_z
for k = 1, 2, ..., N_L do
    for i = 1, 2, ..., N_B do
        Compute x_i = (ξ_{i+1} − ξ_i)/2
        for p = 1, 2, ..., N_GL do
            Compute: β_{i,p} = (ξ_{i+1} − ξ_i) r_p/2 + (ξ_{i+1} + ξ_i)/2
            Assign: X[i,p,k] = x_i, B[i,p,k] = β_{i,p}, W[i,p,k] = w_p
Running in parallel for all r and t
for k = 1, 2, ..., N_L do
    Compute: e_k = η_k/t
    Compute: g_k = γ_k/t
    for i = 1, 2, ..., N_B do
        for p = 1, 2, ..., N_GL do
            Assign: E[i,p,k] = e_k, J[i,p,k] = J_m(r B[i,p,k])
            Generate: the model matrix 𝒦(B[i,p,k], g_k) and the total stress vector Ṽ(B[i,p,k], g_k)
            Compute: Λ̃ = 𝒦(B[i,p,k], g_k)^{-1} Ṽ(B[i,p,k], g_k)
            Extract: ũ_z^{(0)}(B[i,p,k], z, g_k) and Θ̃^{(0)}(B[i,p,k], z, g_k) from the state vector Λ̃
            Assign: U_z[i,p,k] = ũ_z^{(0)}, T_z[i,p,k] = Θ̃^{(0)}
Compute: u_z(r,z,t) = Σ_{i,p,k} E ⊙ X ⊙ W ⊙ J ⊙ B ⊙ U_z
Compute: Θ(r,z,t) = Σ_{i,p,k} E ⊙ X ⊙ W ⊙ J ⊙ B ⊙ T_z
```

Note that $\odot$ denotes the Hadamard product.

**Surface heat source** In reality, the absorbed laser intensity exhibits a continuous exponential decrease along the layer depth according to the Beer-Lambert law:

$$q_e(z) = \frac{P_0}{\pi R^2}\mu_a e^{-(\mu_a+\mu_s')z}, \qquad [106]$$

where $P_0$ is the incident laser power at the absorbing layer plane, $a$ is the laser radius, and $\mu_s' = \mu_s(1-g)$ is the reduced scattering coefficient with $\mu_s$ being the scattering coefficient and $g$ the anisotropy factor. However, based on the construction of the model, the elements of the stress vector correspond to external heat stresses only at interfaces, hence, the $N$-layer analytical model only accepts surface heat sources and not volumetric sources. This limitation is overcome by discretizing the volumetric heat source into planes that are sufficiently close to each other, such that the surface heat source and volumetric heat source become equivalent after a characteristic time that is shorter than the experimental characteristic time (1 ms). Heat diffuses over a thickness of 1 µm with a characteristic time of 2.5 µs. Hence, the volumetric heat deposited on the apical membrane of the RPE (modeled as a layer of 1 µm) can be approximated as a surface heat source, however, the pigmented choroid with a thickness of 20 µm is segmented into five sub-layers of the same thickness with surfacic heat sources placed on top of each sub-layer (7). Consider an absorbing layer of thickness $\ell$ with the top interface at $z = z_H$, the amplitude of the surface heat source was set to equal the total heat deposited over $\ell$, i.e.,

$$q_e(z_H) = \frac{P_0}{\pi R^2}\frac{\mu_a}{\mu_a+\mu_s'}\left(1 - e^{-(\mu_a+\mu_s')\ell}\right). \qquad [107]$$

The complete spatial-temporal profile of the surface heat source is

$$Q_e(r, z_H, t) = q_e(z_H) \times \phi_e(r,t), \qquad [108]$$

where $\phi_e(r,t)$ can be decomposed into spatial and temporal components:

$$\phi_e(r,t) = \phi_{\text{spatial}}(r) \times \phi_{\text{temporal}}(t). \qquad [109]$$

The spatial profile is modeled as

$$\phi_{\text{spatial}}(r) = H(R-r), \qquad [110]$$

where $H$ is the Heaviside function. As shown in Fig. S2B, the measured heating beam at the detector using a model eye lens with 30 mm focal length is the image of a pinhole and its edges are relatively straight. The measured temporal profile was fit to a double exponential function, i.e.,

$$\phi_{\text{temporal}}(t) = [H(t) - H(t-t_0)]\frac{A_1 e^{-t/t_1} + A_2 e^{-t/t_2}}{A_1 + A_2}, \qquad [111]$$

where $t_0 = 10$ ms is the laser pulse duration, $A_1 = 4.63$, $A_2 = 1.01$, $t_1 = 0.35$ ms and $t_2 = 4.98$ ms were fitting results (7).

**Table S1. Initial thermo-mechanical properties for LE retinal layers. $\ell$: layer thickness. $E$: Young's modulus. $\nu$: Poisson's ratio. $\kappa$: coefficient of thermal conductivity. $c_p$: specific heat capacity. $\alpha_{\mathsf{TE}}$: coefficient of thermal expansion. $\mu_a$: absorption coefficient. $\mu_s$: scattering coefficient. $g$: anisotropy.**

| Layer | $\ell$ ($\mu$m) | $E$ (kPa) | $\nu$ | $\kappa$ (W/m·K) | $c_p$ (J/kg·K) | $\alpha_{\mathsf{TE}}$ (1/K) | $\mu_a$ (1/cm) | $\mu_s$ (1/cm) | $g$ |
|---|---|---|---|---|---|---|---|---|---|
| 1. Nerve fiber layer (NFL) | 10 | $1.3^a$ | $0.47^c$ | $0.45^d$ | $4190^f$ | $2.25\times10^{-4}$ | - | - | - |
| 2. Inner plexiform layer (IPL) | 50 | $1.3^a$ | $0.47^c$ | $0.45^d$ | $4190^f$ | $2.25\times10^{-4}$ | - | - | - |
| 3. Inner nuclear layer (INL) | 20 | $2.7^a$ | $0.47^c$ | $0.45^d$ | $4190^f$ | $2.25\times10^{-4}$ | - | - | - |
| 4. Outer plexiform layer (OPL) | 15 | $8^a$ | $0.47^c$ | $0.45^d$ | $4190^f$ | $2.25\times10^{-4}$ | - | - | - |
| 5. Outer nuclear layer (ONL) | 50 | $2.7^a$ | $0.47^c$ | $0.45^d$ | $4190^f$ | $2.25\times10^{-4}$ | - | - | - |
| 6. Connecting cilium layer (CCL) | 10 | 0.02 | $0.47^c$ | $0.45^d$ | $4190^f$ | $2.25\times10^{-4}$ | - | - | - |
| 7. Rod outer segment (ROS) | 24 | $26^a$ | $0.47^c$ | $0.45^d$ | $4190^f$ | $2.25\times10^{-4}$ | - | - | - |
| 8. RPE apical layer (RPEa) | 1 | $5^a$ | $0.47^c$ | $0.45^d$ | $4178^f$ | $2.25\times10^{-4}$ | $9.976\times10^{3\ f}$ | $6.75\times10^{3\ f}$ | $0.84^f$ |
| 9. RPE basal layer (RPEb) | 3 | $5^a$ | $0.47^c$ | $0.45^d$ | $4178^f$ | $2.25\times10^{-4}$ | - | - | - |
| 10. Pigmented choroid layer (PCL) | 20 | $30^a$ | $0.47^c$ | $0.45^d$ | $3840^f$ | $2.25\times10^{-4}$ | $2.494\times10^{3\ f}$ | $2.4\times10^{3\ f}$ | $0.94^f$ |
| 11. Non-pigmented choroid (NPC) | 30 | $30^a$ | $0.47^c$ | $0.45^d$ | $3840^f$ | $2.25\times10^{-4}$ | - | - | - |
| 12. Sclera (SCL) | 500 | $30^a$ | $0.47^c$ | $0.45^d$ | $4178^f$ | $2.25\times10^{-4}$ | - | - | - |
| 13. Fat | $\infty$ | $2^b$ | $0.49^b$ | $0.2^e$ | $2300^b$ | $2.25\times10^{-4}$ | - | - | - |

a. Qu et al. (8). b. Gonzalez et al. (9). c. Lu et al. (10). d. Harting et al. (11). e. Lagendijk (12). f. Goetz et al. (13). The value for the coefficient of thermal expansion is taken from our previous work (7).

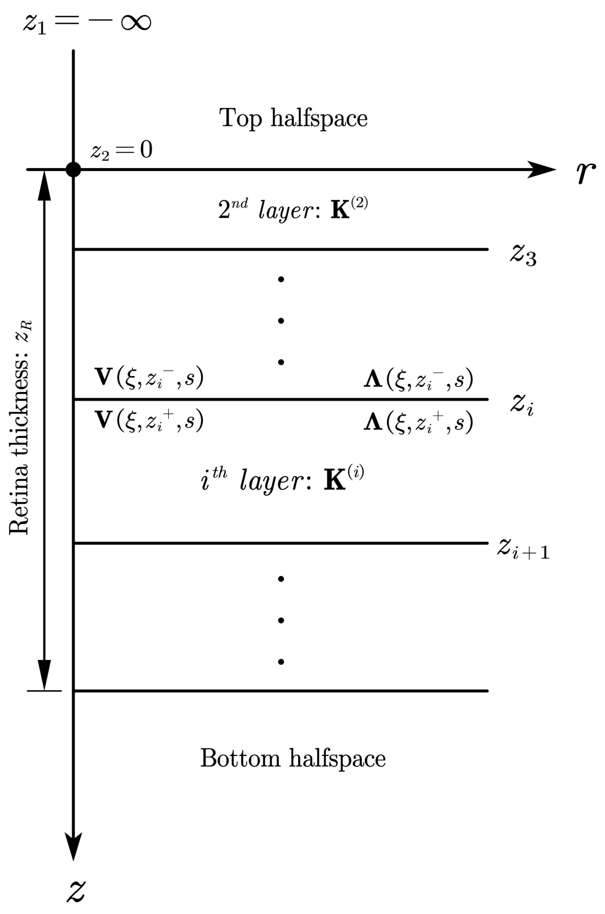


**Fig. S1.** Axisymmetric multilayer model with state vectors, stress vectors, and stiffness matrices at natural layer interfaces.

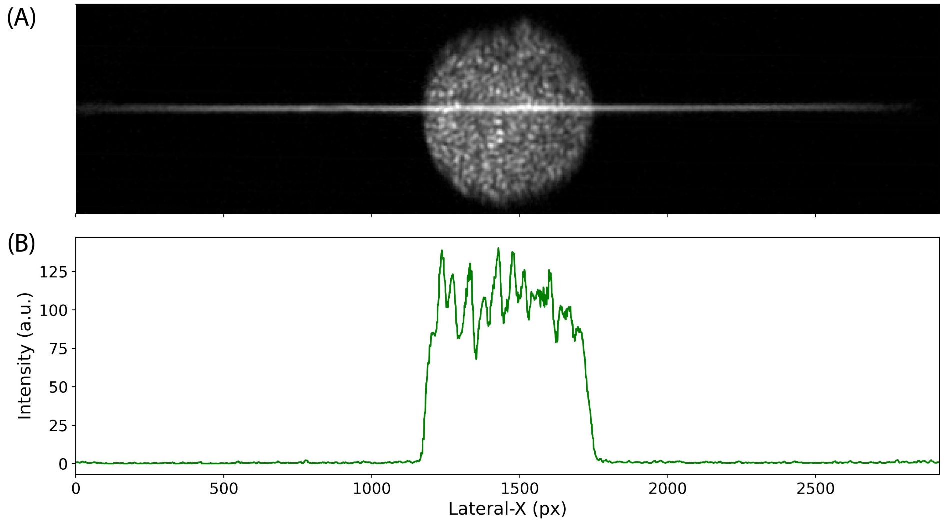


**Fig. S2.** (A) Heating beam and OCT beam at the retinal plane imaged using a model eye lens with a focal length of 30 mm. (B) Heating beam lateral profile. Intensity fluctuations are due to laser speckles.

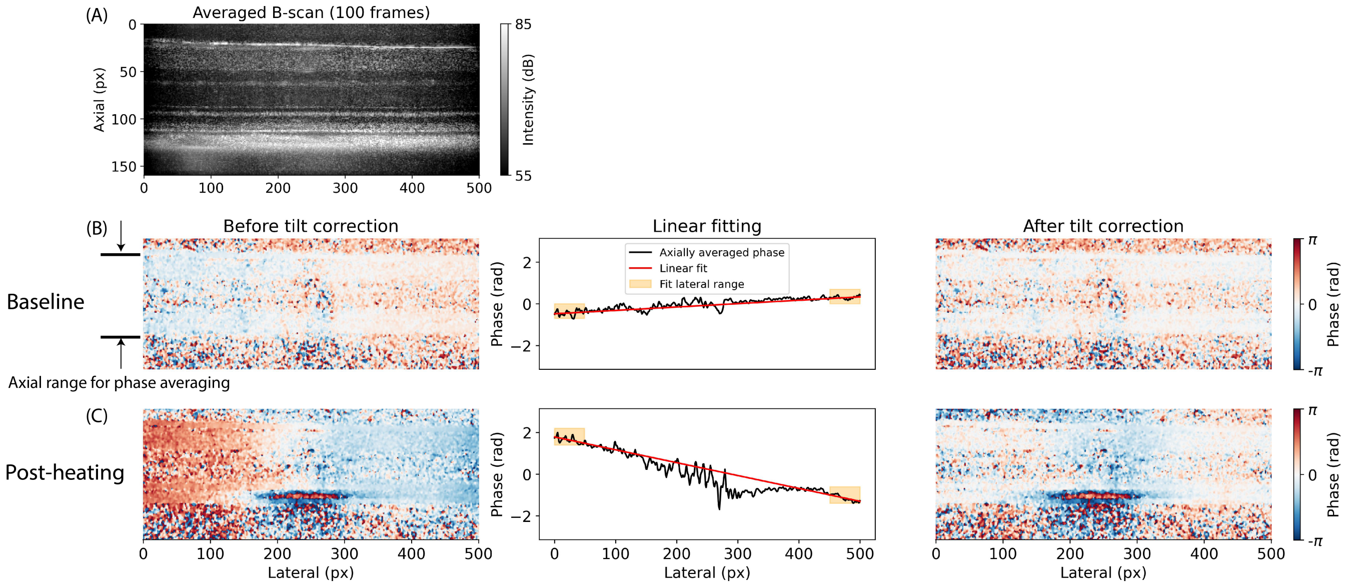


**Fig. S3.** Lateral referencing (the tilt correction). (A) Averaged OCT B-scan of a WT retina (Rat LE 1 in the main text). (B) Baseline measurements (no heating). Raw phase map extracted by time referencing to the first frame (left). The complex valued signals along an A-line were averaged within an axial range to obtain an averaged phase. A linear function is fitted to 50 data points on both ends of the phase profile (center). Each A-line phase was then compensated according to the fitted function value at the corresponding lateral positions (right). (C) Post-heating results.

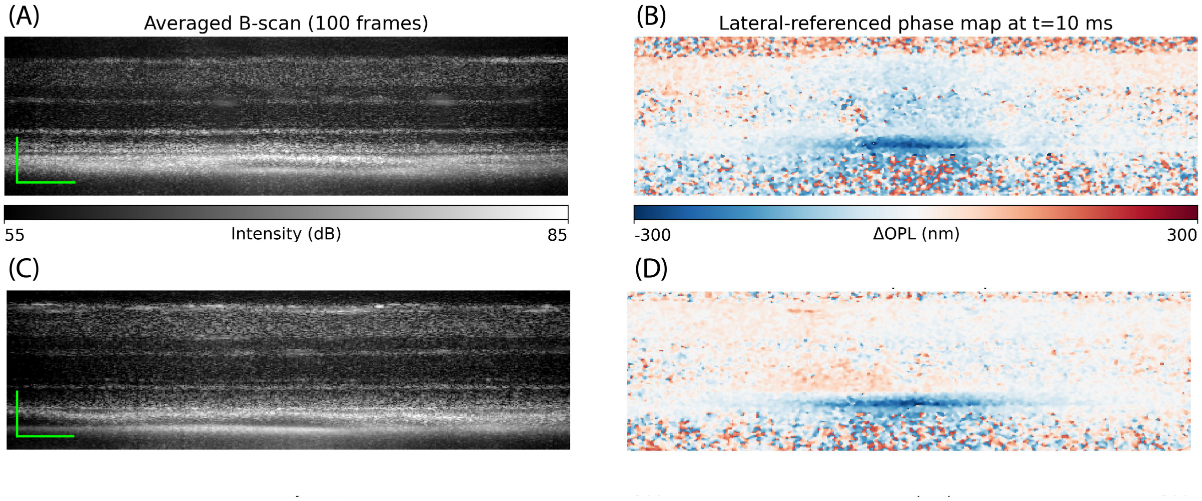


**Fig. S4.** (A-B) Structural image (left) and lateral-referenced phase map (right) for WT animal 4. (C-D) Same for another WT animal. Scale bar: axial, 90 µm; lateral, 50 µm.

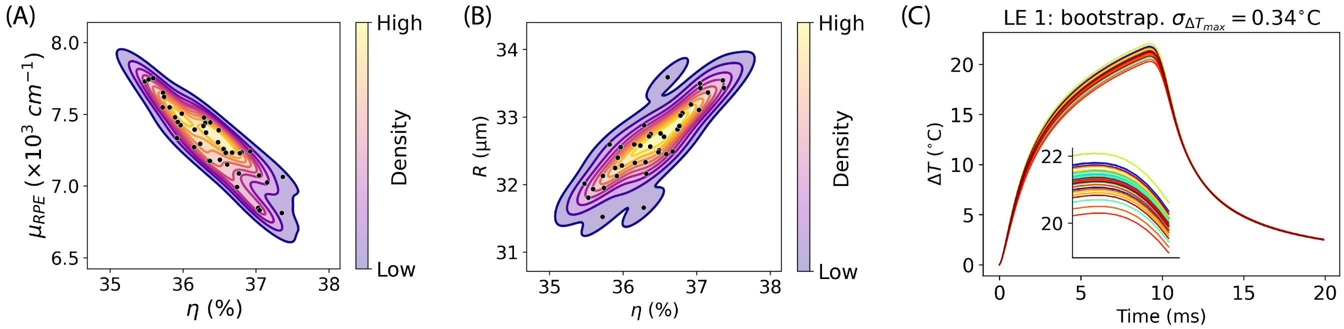


**Fig. S5.** Bootstrapping analysis. (A) Contour plot showing parameter correlations between the absorption coefficient and optical transmittance. Same as Fig. 4E in the main text. (B) Contour plot showing parameter correlations between the heating spot radius and optical transmittance. Naturally, for the same amount of absorbed energy, a smaller spot results in higher temperature, hence also correlated with optical transmittance and absorption coefficient. However, since the laser spot radius also results in faster heat diffusion, and it was mainly fitted through the temporal profile and the lateral extent of $\Delta$OPL, hence, relatively independent from $\eta \times C_\mu$. (C) All predicted temperature profiles from bootstrapping analysis for animal LE 1. Peak temperature standard deviation: 0.34°C.

**Movie S1. Lateral-referenced OPL change for RCS rat. Scale bar: axial, 70 µm; lateral, 50 µm**

**Movie S2. Lateral-referenced OPL change for WT rat. Scale bar: axial, 70 µm; lateral, 50 µm**
